# Recognition of RNA secondary structures with a programmable peptide nucleic acid-based platform

**DOI:** 10.1101/2024.05.14.594238

**Authors:** Rongguang Lu, Liping Deng, Yun Lian, Xin Ke, Lixia Yang, Kun Xi, Alan Ann Lerk Ong, Yanyu Chen, Hanting Zhou, Zhenyu Meng, Ruiyu Lin, Shijian Fan, Yining Liu, Desiree-Faye Kaixin Toh, Xuan Zhan, Manchugondanahalli S. Krishna, Kiran M. Patil, Yunpeng Lu, Zheng Liu, Lizhe Zhu, Hongwei Wang, Guobao Li, Gang Chen

## Abstract

RNA secondary structures comprise double-stranded (ds) and single-stranded (ss) regions. Antisense peptide nucleic acids (asPNAs) enable the targeting of ssRNAs and weakly formed dsRNAs. Nucleobase-modified dsRNA-binding PNAs (dbPNAs) allow for targeting of relatively stable dsRNAs. A programmable RNA structure-specific targeting strategy is needed for simultaneous recognition of dsRNAs and ssRNAs. Here, we report on combining dbPNAs and asPNAs (designated as daPNAs) for the targeting of dsRNA-ssRNA junctions. Our binding and modeling data suggest that combining traditional asPNA (with a 4-letter code: T, C, A, and G) and dbPNA (with a 4-letter code: T or s^2^U, L, Q, and E) scaffolds facilitates RNA structure-specific tight binding (nM to μM) under physiologically-relevant conditions. We further applied our daPNAs in substrate specific inhibition of Dicer acting on pre-miR-198 in a cell-free assay and regulating ribosomal frameshifting induced by model hairpins in both cell-free and cell culture assays. daPNAs would be a useful platform for developing chemical probes and therapeutic ligands targeting RNA.

**Highlight:**

- We demonstrated that sequence- and structure-specific targeting of RNA can be facilitated by nucleobase-modified dsRNA-binding PNAs (dbPNAs) platform in combination with antisense PNAs (asPNAs). We name the novel PNAs as daPNAs.
- daPNAs can be used in a programmable way for targeting RNAs by formation of a short triplex next to a short duplex at a dsRNA-ssRNA junction.
- We applied our daPNAs in substrate specific inhibition of Dicer acting on pre-miR-198 in a cell-free assay and regulating ribosomal frameshifting induced by model hairpins in both cell-free and cell culture assays.
- The daPNAs platform would serve as useful junction-specific molecular glues for the targeting of many biologically important RNA structures in transcriptomes.

## INTRODUCTION

RNAs perform diverse catalytic and regulatory functions in viruses and cells. RNA structures, critical for RNA functions, are mainly stabilized by base-paired double-stranded (ds) stem regions (1, 2). Together with single-stranded (ss) loop regions, RNAs can fold into complex secondary and tertiary structures,(3) facilitating the molecular recognition of RNA structures by small molecules (4-7) and peptides/proteins (8, 9). Antisense oligonucleotides (ASOs) are a successful class of RNA-targeting drugs which bind to target RNAs by Watson-Crick base pairing. ASOs have already been approved for the treatment of certain diseases, such as Duchenne Muscular Dystrophy (DMD) and Spinal Muscular Atrophy (SMA) (10). ASOs typically range in size from 13-25 nucleotides (nt) (10).

Currently, programmable RNA structure-specific and tight-binding ligands are relatively unexplored. Peptide nucleic acid (PNA) is characterized by a neutral, peptide-like backbone, which imparts chemical stability and nuclease resistance **(Figure 1)** (11). PNA-DNA and PNA-RNA duplexes are generally more stable than DNA-DNA and RNA-RNA duplexes (12-14). Strong binding of a PNA to a complementary sequence allows the strand invasion of a dsRNA or a dsDNA, with one of the strands of the dsRNA/dsDNA displaced by the PNA. Importantly, PNAs show promising bioactivities in animal models (15-22). However, traditional antisense PNAs (asPNAs) do not bind to RNAs by recognizing RNA structures, which may give rise to off-target binding.

**Figure 1.**
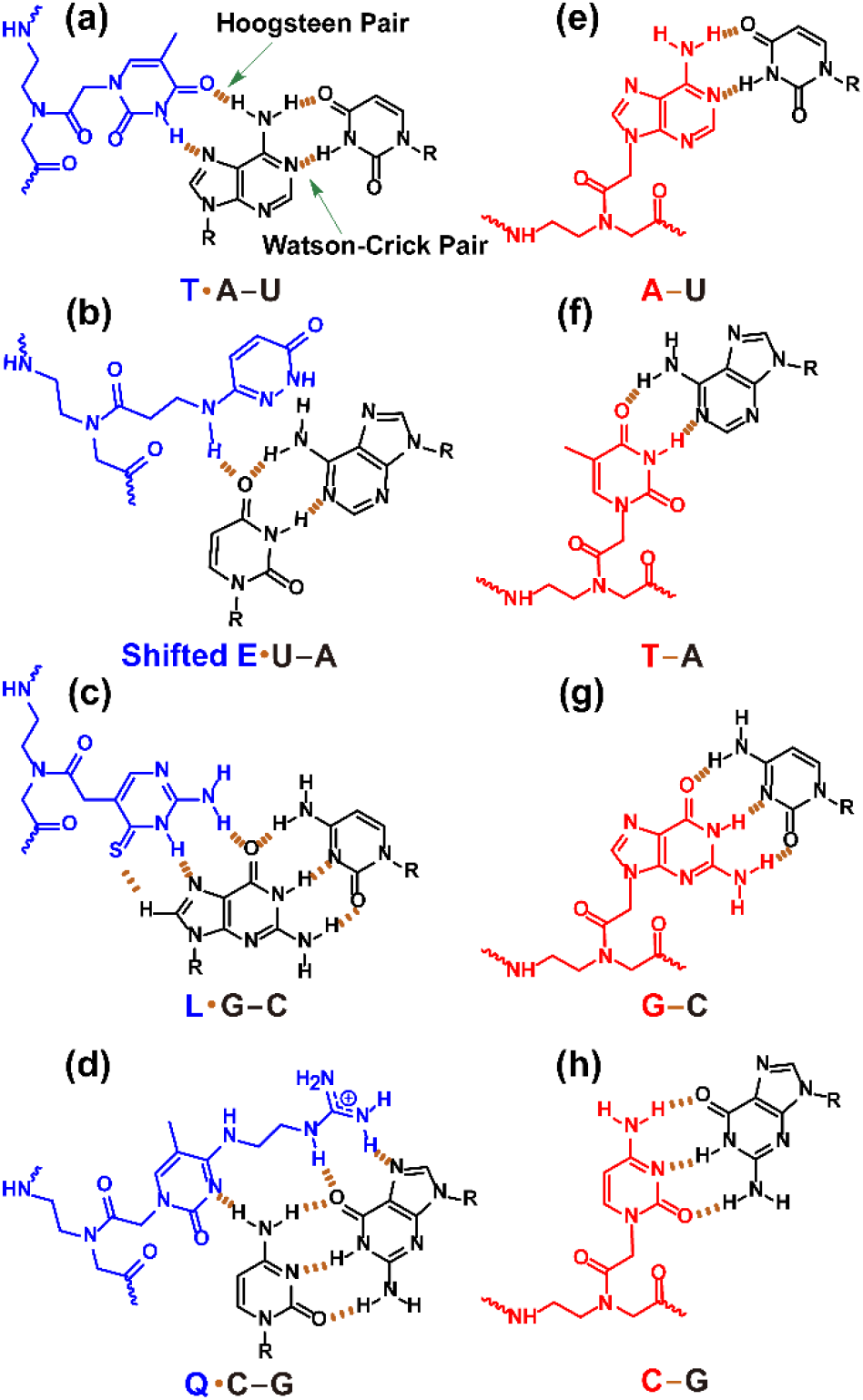
Chemical structures of base triples and base pairs formed between PNA and RNA. “R” represents the sugar-phosphate backbone of RNA. Hydrogen bonding interactions are indicated by orange dashed lines. (a-d) Base triples of T·A-U, Shifted E·U-A, L·G-C, Q·C-G**(28)** formed by dbPNA residues with RNA base pairs. Compared to unmodified C base, modified L base shows enhanced recognition of a G-C pair with reduced pH dependence**(23)**. The enhanced van der Waals interaction between the sulfur and hydrogen atoms is shown. The Hoogsteen and Watson-Crick pairs are shown as dots and lines, respectively. PNA and RNA are shown in blue and black, respectively. The base triple formation allows for the recognition of dsRNA regions. (e-h) Base pairs of A-U, T-A, G-C and C-G formed by asPNA with ssRNA. Unmodified PNA and RNA are shown in red and black, respectively. The base pairing between PNA and RNA allows the recognition of ssRNA regions.

We and others have shown that short (e.g., 10 mer) dsRNA-binding PNAs (dbPNAs) incorporated with modified nucleobases can form sequence-specific triplexes with dsRNAs through the formation of consecutive PNA·RNA-RNA base triples **(Figure 1a-d)**, with significantly weakened binding to dsDNAs and ssRNAs (23-35). The dbPNAs have shown biological activities in stimulating ribosomal frameshifting (36), and inhibiting translation (37), miRNA maturation (27, 38), RNA editing (39), and viral RNA replication (26). However, naturally occurring dsRNAs in functional RNAs are often interrupted by ssRNA loop regions, which may limit the application of dbPNAs. Thus, developing a scaffold for the simultaneous recognition of dsRNA and ssRNA regions is critical. We herein designed novel PNAs combining dbPNAs (containing bases of T or s^2^U, L, Q, and E or S, **Figures 1a-d, S1**) (23, 24, 26, 27, 40) and traditional asPNAs (containing unmodified bases T, C, G, and A, **Figure 1e-h**) (designated as daPNAs, see **Figure 2**) for the simultaneous recognition of dsRNA and immediately adjacent ssRNA regions. Binding data suggest that novel daPNAs can be utilized for targeting the dsRNA-ssRNA regions, including those found in model RNA hairpins, microRNA precursor, and tau pre-mRNA splice site hairpin. Our cell-free and cell-based assays show that daPNAs exhibit biological activities. daPNAs can be utilized to target dsRNA-ssRNA regions with high sequence/structure specificity and strong binding affinity, potentially overcoming the obstacles encountered by other techniques, such as small molecules and traditional ASOs, including asPNAs.

**Figure 2.**
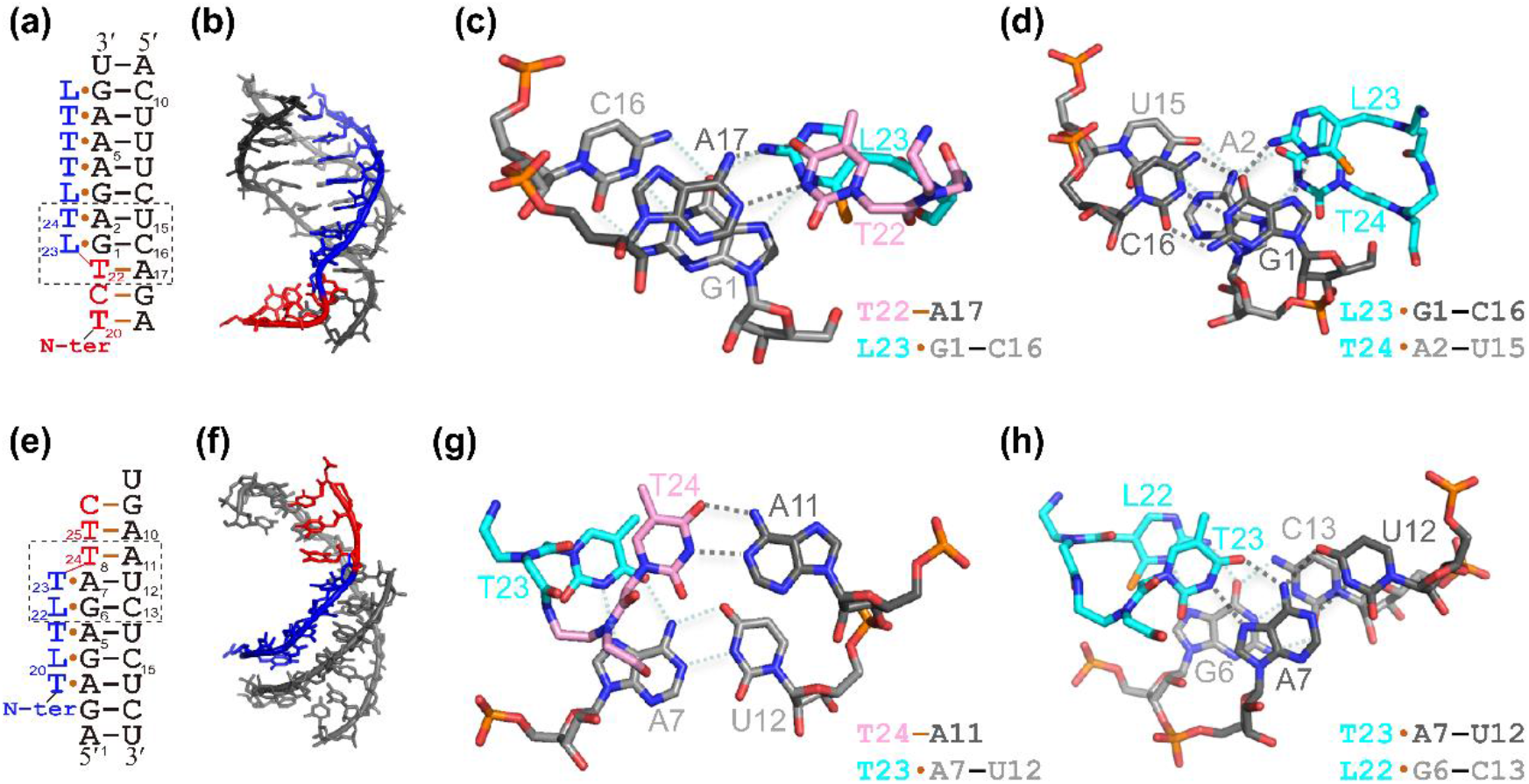
Modelled structures of daPNA-RNA complexes. (a-d) A modelled daPNA·(dsRNA-ssRNA) structure with the N-terminal segment of the PNA forming a PNA-RNA duplex. (a,b) Secondary structure scheme and modeled structure, with the RNA and PNA residues shown in black and color, respectively. The detailed stacking patterns in the triplex-duplex junctions are shown in panels c and d, with the carbon atoms for the bases of RNAs and PNAs shown in black and color, respectively. (e-h) A modelled daPNA·(dsRNA-ssRNA) structure with the C-terminal segment of the PNA forming a PNA-RNA duplex. (e,f) Secondary structure scheme and modeled structure, with the RNA and PNA residues shown in black and color, respectively. The detailed stacking patterns in the triplex-duplex junctions are shown in panels g and h, with the carbon atoms for the bases of RNAs and PNAs shown in black and color, respectively.

## RESULTS AND DISCUSSION

### daPNA binding to model dsRNA-ssRNA junction and stimulate ribosomal frameshifting

We designed a series of short daPNAs with the asPNA segment antiparallel to the ssRNA region and the dbPNA segment forming a parallel triplex with the dsRNA region (see **Figure 2a,e** for example, see all the PNAs studied in **Table S1 and Figures S2-S5**). Our structural modeling study following the reported method(25, 41) suggests that such a novel fold with a short major-groove dbPNA·dsRNA triplex next to a short asPNA-ssRNA duplex at a dsRNA-ssRNA junction is structurally compatible without significant distortion in the backbones and bases **(Figures 2b,f, S6 and S7)**. The detailed structures of the dsRNA-ssRNA junctions in complex with daPNAs are shown in **Figure 2c,d,g,h**. Importantly, the junction structures complexed with daPNAs are largely stable with a force field (42) in the modeling even without base-base hydrogen bonding restraints, although local dynamics are observed (**Figure S7**).

We constructed a hairpin rHP1-5t containing a 5-nucleotide (nt) 3′ dangling end (overhang) as a model of dsRNA-ssRNA motif (**Figure 3a**). We made a 10-mer daPNA da5t (NH_2_-Lys-AGAGTLTLTT-CONH_2_) targeting rHP1-5t (**Figure 3a,b**). The 10-mer daPNA da5t contains an N-terminal 4-mer segment (AGAG) for the formation of an antiparallel PNA-RNA duplex with the 3′ dangling end and a C-terminal 6-mer dbPNA segment (TLTLTT) for the formation of a major-groove PNA·dsRNA triplex (**Figures 2 and 3a**). We carried out non-denaturing polyacrylamide gel electrophoresis (PAGE) and bio-layer interferometry (BLI) to characterize the recognition of the model dsRNA-ssRNA junction by PNAs (**Figure 3c,d**). Non-denaturing PAGE data suggest that da5t indeed binds tightly to rHP1-5t (*K*_D_=4±1 nM) **(Figures 3c**,**e, S8)**. However, da5t shows no binding with rHP1-3t, with a 5′ dangling end with the same ssRNA sequence as rHP1-5t **(Figures 3a,e, S8)**. Consistent with the PAGE results, bio-layer interferometry (BLI) data demonstrated that da5t can bind to rHP1-5t with a *K*_D_ of 7±10 nM but with no significant binding to rHP1-3t **(Figures 3d,e, S9)**. The data suggest that the asPNA segment of a daPNA forms an antiparallel duplex but not parallel duplex with ssRNA in a dsRNA-ssRNA junction. Furthermore, da5t does not bind with rHP1, which has no ssRNA dangling end, as determined by PAGE and BLI assay **(Figures 3e, S8, S9)**. Taken together, our data suggest that daPNA binds to dsRNA-ssRNA junction structure with structure specificity. We subsequently generated three rHP1-5t mutants (rHP1-5t-m1, rHP1-5t-m2 and rHP1-5t-m3) with U29C, A6C&U27G and G7C&C26G mutations, respectively (**Figure 3a)**, to test the sequence specificity of da5t. Both the PAGE and BLI results revealed that da5t has a weakened binding to rHP1-5t-m1 **(Figures 3e, S8, S9)**, indicating that the asPNA segment of da5t is an important part of a daPNA in binding with an RNA structure. The rHP1-5t-m2 and rHP1-5t-m3 have the single base pair mutation at the junction and adjacent to the dsRNA-ssRNA junction, respectively **(Figure 3a)**. da5t shows significantly weakened or no binding to rHP1-5t-m2 and rHP1-5t-m3 **(Figures 3e, S8, S9)**, suggesting that an internal or terminal base triple is critical for da5t binding. A daPNA negative control (da5t-NC, NH_2_-Lys-TGTATAQLL-CONH_2_) shows no observable binding with rHP1-5t **(Figure S8)**. We previously reported that a dbPNA (P5) forms a triplex with rHP1 (**Figure 3a**) with a *K*_D_ of about 200 nM (23, 36). Compared to da5t, dbPNA P5 binds to rHP1, rHP1-5t, and rHP1-3t with a similar but comparably weakened binding affinity **(Figure 3e, S8, S9)** (36).

**Figure 3.**
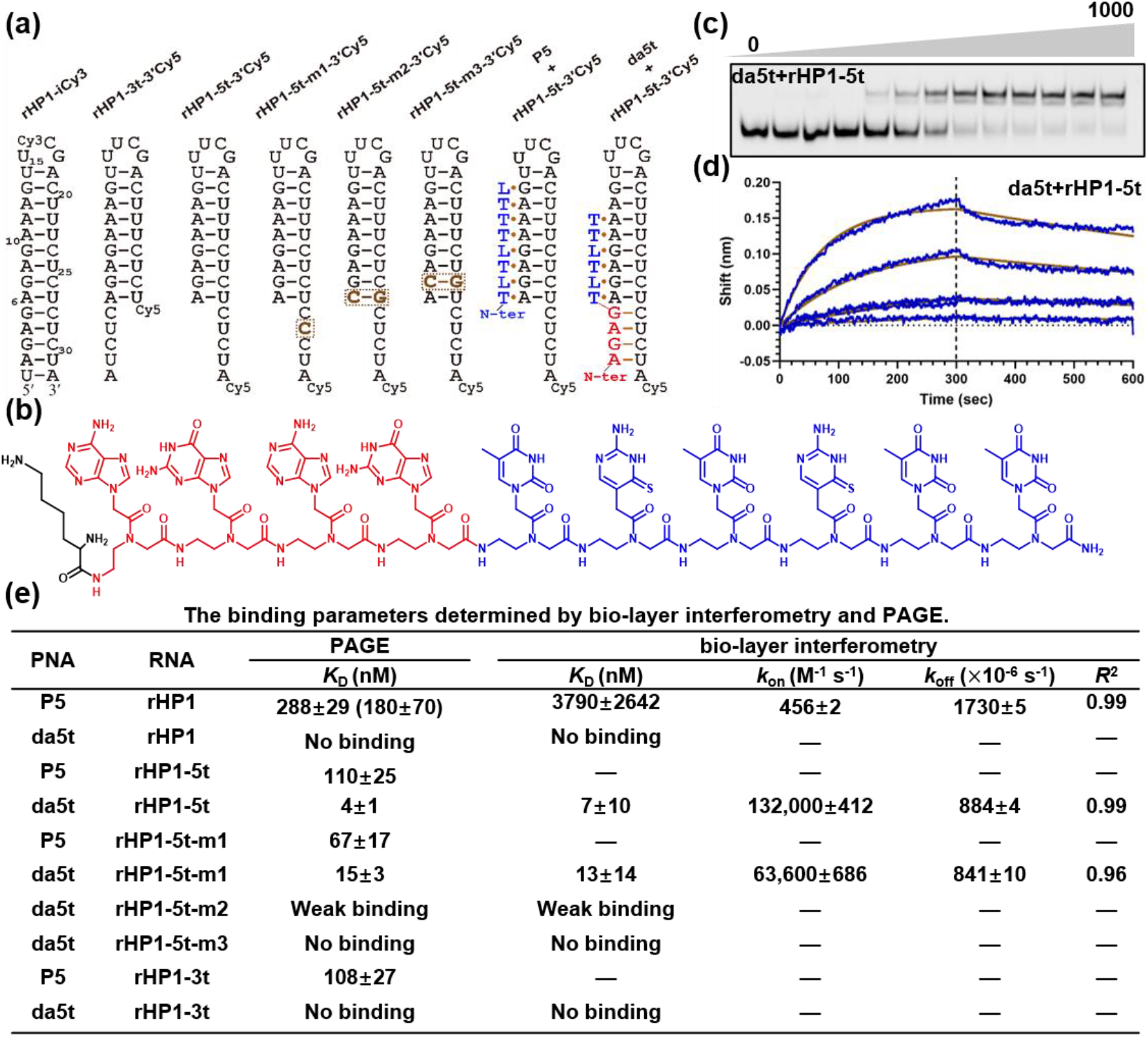
Targeting model RNAs with dsRNA-ssRNA junctions. (a) RNA constructs and schematics of a dbPNA and a daPNA binding to rHP1-5t. (b) Chemical structure of daPNA da5t (NH_2_-Lys-AGAGTLTLTT-CONH_2_). The dbPNA and asPNA segments are shown in blue and red, respectively. The lysine residue (black) is in L configuration. (c, d) Representative binding data of PNA da5t with rHP1-5t base on nondenaturing PAGE and bio-layer interferometry assay, respectively. (e) Binding parameters obtained by nondenaturing PAGE and BLI. The data shown in the parentheses is from ref(27).

We further applied daPNAs in targeting mRNA structures for translational regulation. Programmed −1 ribosomal frameshifting (PRF) is a recoding mechanism utilized by many RNA viruses to express viral proteins in the −1 frame and 0 frame with defined ratios. The mRNA elements important for PRF include a slippery sequence (e.g., U UUU UUA in 0 frame and UUU UUU A in −1 frame), single-stranded spacer (about 8 nt), and downstream mRNA structure **(Figure 4a-c)** (43-48). Dual luciferase reporter assay **(Figure 4a)** is a documented method for studying PRF (36, 49, 50). We previously reported that dbPNA P5 **(Figure 3a)** forms a triplex with rHP1 and stimulates PRF efficiency by enhancing the stability of rHP1 at the ribosomal entry site (23, 36).

**Figure 4.**
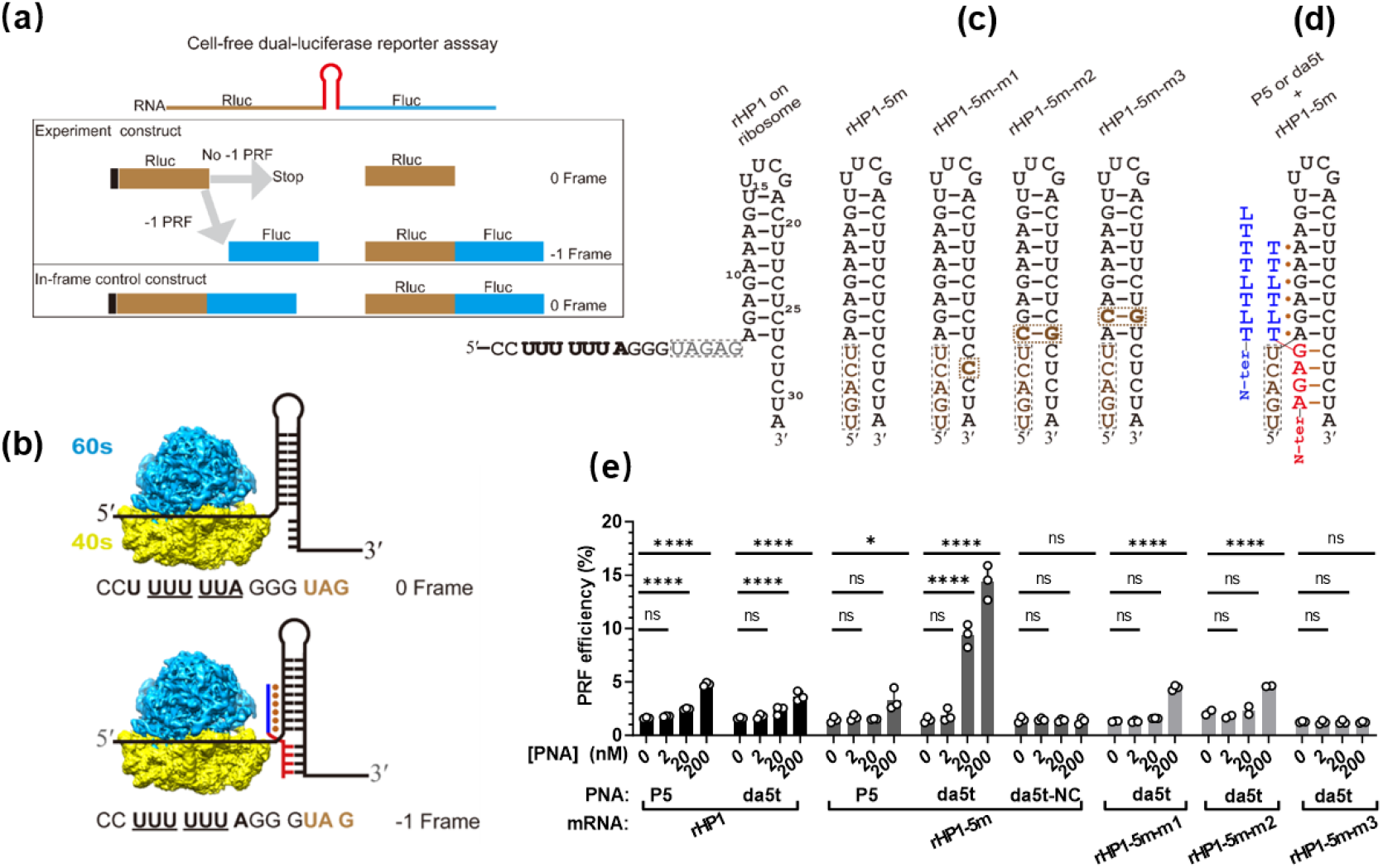
Schematic of dual-luciferase reporter system for measuring programmed −1 ribosomal frameshifting (PRF) efficiency and cell-free dual-luciferase reporter assay data for dbPNA and daPNA targeting model RNA hairpins. (a) Schematic of the dual-luciferase reporter to determine the PRF efficiency in the cell-free system. (b) Ribosome positioned at slippery site with partially open rHP1 RNA structure at the mRNA entry site. A −1 ribosomal frameshifting results in the reading frame shifted from 0 frame (U UUU UUA) to −1 frame (UUU UUU A), causing the read through of the stop codon UAG (shown in brown) at the 0 frame. daPNA binding at the dsRNA-ssRNA junction at the mRNA entry site results in an enhanced −1 ribosomal frameshifting. (c) Secondary structures of rHP1 with the bottom 5 base pairs unwound by ribosome positioned at the slippery site and rHP1-5m and mutants. The mutated residues are shown in brown. (d) Schematic of dbPNA P5 and daPNA da5t binding to rHP1-5m. (e) Cell-free dual-luciferase reporter assay data for dbPNA and daPNA targeting model RNA hairpins. The data were analyzed by GraphPad Prism 9.3 and calculated by an ordinary one-way analysis of variance (ANOVA) using Dunnett’s multiple comparisons test against the mean of mRNA alone group. The error bars represent ± S.D. * P<0.05, ** P<0.01, *** P< 0.001, **** P<0.0001, ns: not significant.

Herein, we employed dual-luciferase reporter assay to test if daPNA binding to mRNA structures affect PRF. We hypothesize that daPNA binding to mRNA structure at the ribosome entry site with ribosome positioned at the slippery site (**Figure 4b,d**) may temporarily inhibit elongating ribosome’s helicase activity in unwinding mRNA secondary structure (51, 52). With ribosome positioned at the slippery site, it is expected that the bottom five base pairs of rHP1 are unwound, resulting in the formation of a ribosome-induced dsRNA-ssRNA junction identical to that of rHP1-5t **(Figures 3a, 4b,c)**. We also made mRNA constructs rHP1-5m and rHP1-5m-m1/rHP1-5m-m2/rHP1-5m-m3 to mimic the ribosome-independent dsRNA-ssRNA junction structures present in rHP1-5t and rHP1-5t-m1/rHP1-5t-m2/rHP1-5t-m3, respectively **(Figures 3a and 4c)**. The schematic of dual-luciferase reporter plasmid (pDL) is shown in **Figure 4a** with the inserted sequences verified by sequencing **(Table S3, Figure S10a)**. Cell-free dual-luciferase reporter assay data show that rHP1, rHP1-5m, rHP1-5m-m1, rHP1-5m-m2, rHP1-5m-m3 have similar PRF efficiencies of 1.9±0.1%, 1.5±0.2%, 1.3±0.1%, 1.9±0.1% and 1.3±0.1%, respectively. Consistent with our previous report(36), applying 200 nM dbPNA P5 results in moderately enhanced PRF efficiencies of 4.8±0.2% and 3.3±1.0%, for rHP1 and rHP1-5m, respectively **(Figures 4e, S11a**,**b)**.

Consistent with the proposed binding of daPNA da5t to the mRNA dsRNA-ssRNA junction at the ribosome entry site **(Figure 4b,d)**, applying 200 nM da5t causes significantly enhanced PRF efficiency of rHP1-5m (14.8±1.6%) **(Figure 4e)**. da5t binds to rHP1-5m in the absence of an elongating ribosome as the dsRNA-ssRNA junction structure is independent of a ribosome positioned at the slippery site (**Figure 4d**). Interestingly, da5t also stimulates moderately the PRF efficiency of mRNA rHP1 **(Figure 4e)**, even though da5t shows no binding to short free hairpin rHP1 **(Figure 3e)**. It is likely that the ribosome positioned at the slippery site results in the unwinding of bottom 5 base pairs of rHP1 structure in the mRNA (**Figure 4b**,**c**), with the intermediate structure allowing for the binding of da5t resulting in frameshifting stimulation. It’s important to note that applying da5t stimulates the expression of Fluc, but not the Rluc **(Figure S11a**,**b)**. Applying a control PNA da5t-NC at 200 nM shows no observable change in the PRF efficiency of rHP1-5m (**Figures 4e, S11a**,**b**).

rHP1-5m-m1 and rHP1-5m-m2/rHP1-5m-m3 have a single base mutation and a single base pair mutation, respectively, in the ssRNA and dsRNA regions of rHP1-5m **(Figure 4c)**. As expected, rHP1-5m-m1 and rHP1-5m-m2 show a much-reduced response to the application of da5t in stimulating frameshifting, with rHP1-5m-m3 showing no response to the addition of da5t **(Figures 4e, S11a**,**b)**. Taken together, our cell-free dual-luciferase data clearly suggest that da5t binds to the dsRNA-ssRNA junction resulting in the sequence and structure specific enhancement of ribosomal frameshifting.

We further employed a dual-fluorescent reporter (eGFP/mCherry) assay to determine the effect of daPNA on PRF efficiency in cell culture.(8, 44) We constructed a dual-fluorescent reporter plasmid (pDF) derived from pcDNA5 vector, which contains a P2A ribosome skip signal sequence downstream of the frameshifting stimulation elements between eGFP and mCherry coding sequences **(Figure 5a and Table S3)**. Twenty-four hours after the plasmid and PNA transfection into human embryonic kidney 293T (HEK-293T) cell, each cell that shows eGFP fluorescence signal was used for the PRF efficiency analysis **(Figure S12)**. mCherry level shows no correlation with the eGFP level, indicating that the plasmid transfection efficiency does not affect significantly the PRF efficiency **(Figure S12)**. The cell-based assay using pDF reporter system gives a PRF efficiency value of 4.5±0.2% for rHP1-5m **(Figure 5b)**. da5t shows a potent cellular activity in stimulating PRF efficiency in a dose dependent manner, with an efficiency of 9.2±0.9% at 10 μM **(Figure 5b)**. However, applying 10 μM P5 only results in a slight increase of PRF efficiency to 5.6±0.4% **(Figure 5b)**. da5t shows no clearly observable activity in stimulating the PRF of rHP1-5m-m1 **(Figure 5b)**, indicating the cellular activity is sequence specific. Importantly, da5t only stimulates the expression of mCherry, with the eGFP expression unchanged **(Figure S11c**,**d)**. A histogram analysis of the PRF efficiency of individual eGFP-active cells clearly suggests that da5t is a specific stimulator of PRF of rHP1-5m **(Figures 5c, S12)**. Together, our data suggest that daPNA molecules are useful in targeting mRNA structures at the ribosomal entry site for stimulating ribosomal frameshifting.

**Figure 5.**
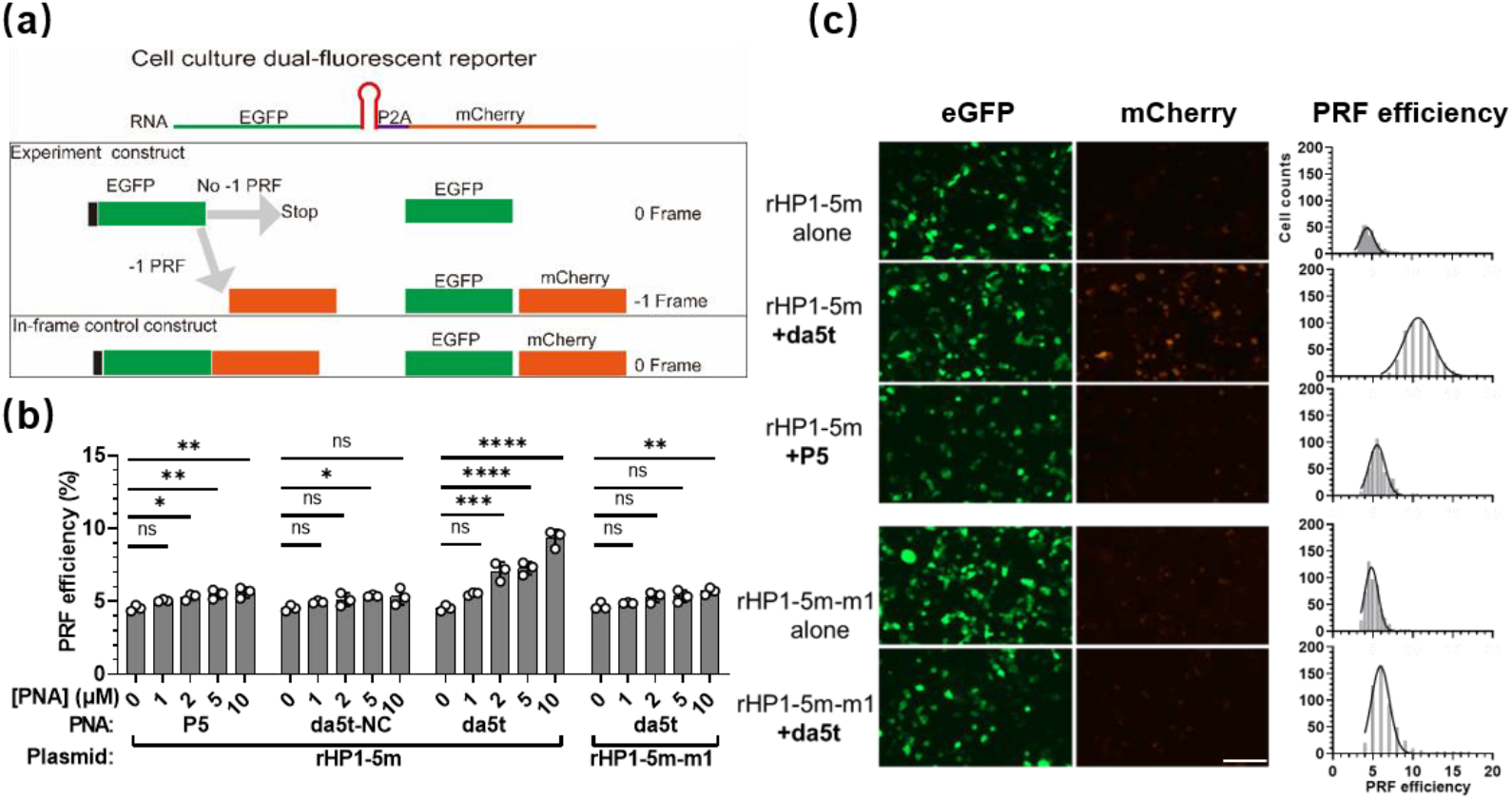
Schematic of dual-fluorescent reporter (eGFP/mCherry) system for measuring programmed −1 ribosomal frameshifting (PRF) efficiency and cell culture dual-fluorescent reporter assay data for dbPNA and daPNA targeting model RNA hairpins. (a) Schematic of the dual-fluorescent reporter. (b) PRF efficiency stimulating by daPNA in cell culture. See the PNA sequences in Figure 4. (c) Representative fluorescence imaging data and histogram analysis of PRF efficiencies of eGFP positive cells with and without 10 μM PNAs. The scale bar is 50 μm. The data were analyzed by GraphPad Prism 9.3 and calculated by an ordinary one-way analysis of variance (ANOVA) using Dunnett’s multiple comparisons test against the mean of plasmid alone group. The error bars represent ± S.D. * P<0.05, ** P<0.01, *** P<0.001, **** P<0.0001, ns: not significant.

### daPNAs targeting pre-miRNA-198 inhibits Dicer cleavage

We next tested the generality of the application of the daPNA platform by designing a daPNA targeting a biomedically important regulatory precursor microRNA (pre-miR) pre-miR-198 **(Figure 6a**) (27, 53). We previously demonstrated that an 8-mer dbPNA, db-198 (NH_2_-Lys-LLTL2TLL-CONH_2_, 2: s^2^U, see **Figure S1**) binds to pre-miR-198 with a *K*_D_=80±50 nM (27). In this study, we made a daPNA by adding three adenine residues at the C-terminal side of db198 (da198-AAA, NH_2_-Lys-LLTL2TLLAAA-CONH_2_) (**Figure 6a**). We characterized the binding properties by nondenaturing PAGE and BLI (**Figure 6b,c**). As expected, the nondenaturing PAGE data show that, compared to the original dbPNA (db198), da198-AAA shows enhanced binding (*K*_D_=10±3 nM) **(Figure 6b,d, S13)**. The BLI results also show that da198-AAA (*K*_D_=278±188 nM) has a stronger binding than db198 (*K*_D_=787±553 nM) with pre-miR-198 **(Figures 6c,d, S14)**.

**Figure 6.**
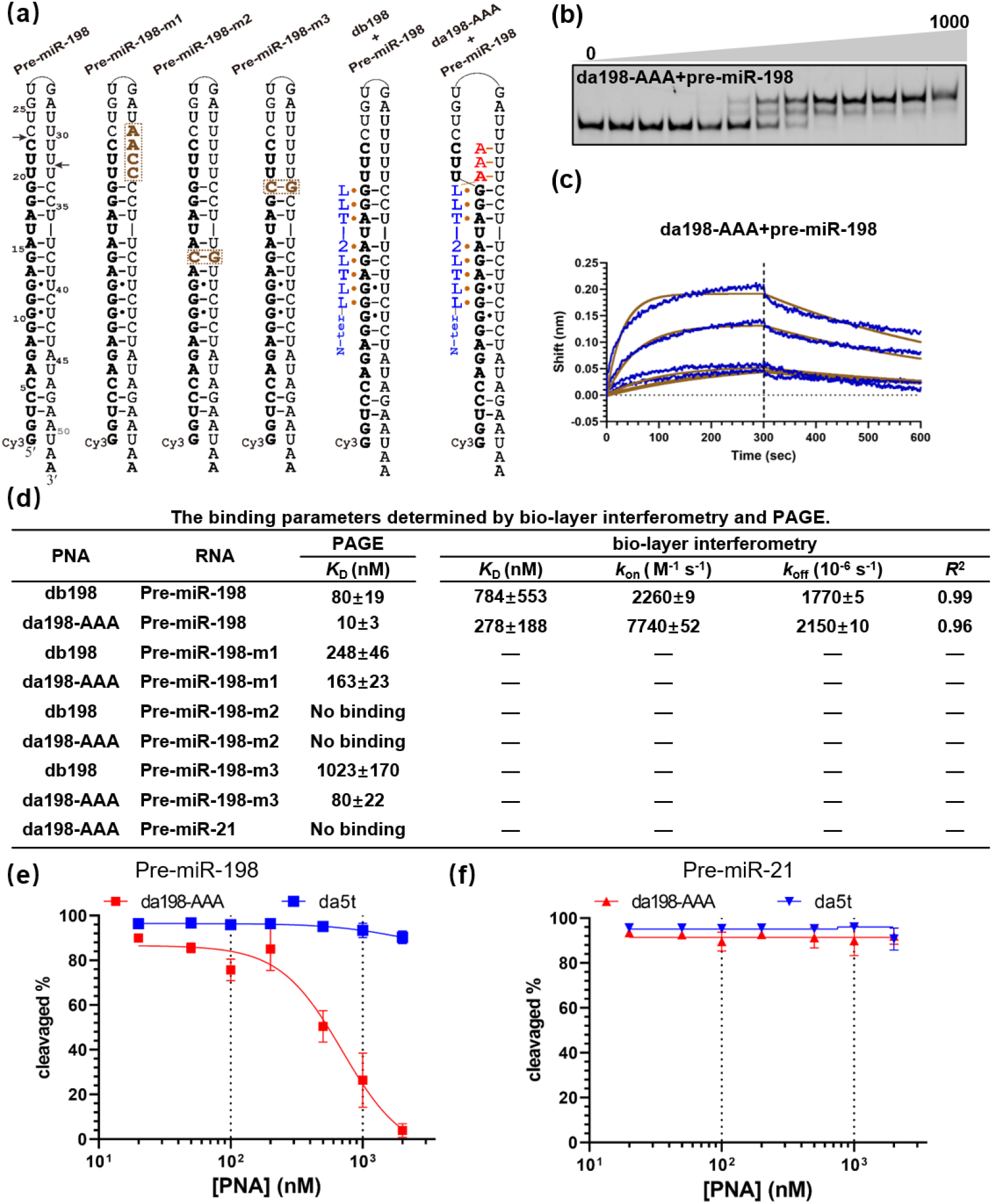
daPNAs binding to pre-miR-198 and inhibition of Dicer enzyme cleavage activity. (a) Schematic of pre-miR-198 hairpin targeting by dbPNA db198 and daPNA da198-AAA. The mature miR-198 sequence is shown in bold. The Dicer cleavage sites for pre-miR-198 are indicated by arrows. The mutated RNA constructs are shown with the mutated residues shown in brown. (b,c) Representative binding data of PNA da198-AAA with pre-miR-198 base on nondenaturing PAGE and bio-layer interferometry assay, respectively. (d) Binding parameters obtained by nondenaturing PAGE and BLI. (e, f) Cell-free Dicing data show that daPNA da198-AAA designed for targeting pre-miR-198 inhibits the Dicer cleavage activity specifically on pre-miR-198.

We subsequently generated three pre-miR-198 mutants (pre-miR-198-m1, pre-miR-198-m2 and pre-miR-198-m3) with residues at 30-33 changed from UUUU to AACC, and with G14C&C38G and G19C&C34G mutations, respectively (**Figure 6a)** to test the sequence specificity of da198-AAA. The nondenaturing PAGE data show that pre-miR-198-m1 binds to da198-AAA with a *K*_D_ of 163±23 nM which is about 16-fold weaker than that of pre-miR-198 (**Figures 6d, S13)**. However, the *K*_D_ of 248±46 nM for pre-miR-198-m1 with db198 is only 3-fold weaker than that of pre-miR-198 (**Figures 6d, S13)**. These results demonstrate that the PNA-RNA duplex formation between the asPNA part of da198-AAA and the loop region U residues of pre-miR-198 significantly enhances the binding affinity. Furthermore, pre-miR-198-m2 shows no binding to both da198-AAA and db198 (**Figures 6d, S13)** suggesting that an internal PNA·RNA-RNA base triple formation is critical for RNA structure recognition. Compared with pre-miR-198, pre-miR-198-m3 shows significantly weakened binding to da198-AAA (*K*_D_=80±22 nM) and db198 (*K*_D_=1023±170 nM) **(Figures 6d, S13)**. The data suggest that a PNA·RNA-RNA base triple formed at the dsRNA-ssRNA junction region is also important for the RNA structure recognition.

We further tested if the newly made db198-AAA can inhibit pre-miR-198 maturation catalyzed by Dicer. The Dicer cleavage assay data show that da198-AAA can specifically inhibit the Dicer cleavage activity on pre-miR-198, but not pre-miR-21 *in vitro* **(Figures 6e,f, S15 and S16)**. Taken together, our data daPNA platform molecules are useful in sequence and structure specifically targeting RNA structures for inhibiting Dicer and other enzymes acting on RNA.

### Applications of daPNAs in binding to other RNAs

Finally, we tested the application of the daPNA platform for targeting tau pre-mRNA splice site hairpin structure **(Figure S17a)**. The tau pre-mRNA hairpin structure **(Figure S17a)** is located at the exon 10-intron 10 junction (54-58). Point mutations in this splice site hairpin structure results in (de)stabilizing of the hairpin and thus aberrant alternative splicing leading to neurodegenerative diseases such as frontotemporal dementia with 15 Parkinsonism linked to chromosome 17 (FTDP-17) (56-61). Significantly, our nondenaturing PAGE result suggests that 9-mer daPNA da-tau (NH_2_-Lys-TQTQTGCCG-CONH_2_) **(Figure S17b)** binds strongly to the tau pre-mRNA splice site hairpin (tau-Cy3) at 200 mM NaCl, pH 7.5 (*K*_D_=27±11 nM, **Figure S17b**). The advantage of a daPNA compared to a traditional asPNA is that daPNA is sequence and structure-specific and has a relatively lower energy barrier for melting the preformed RNA structures. In this case, the relatively weak bottom stem of tau pre-mRNA splice hairpin structure (57, 61) is targeted by strand invasion by a 5-mer asPNA segment of da-tau to form a PNA-RNA duplex, which is presumably cooperatively stabilized by a 4-base-triple PNA·dsRNA triplex.

## Conclusion

In summary, we have developed a novel and generally applicable binding mode by combining dbPNA and asPNA for the recognition of a dsRNA region and an immediately adjacent ssRNA region (dsRNA-ssRNA junction) through the simultaneous formation of an asPNA-RNA duplex and a major-groove dbPNA·RNA_2_ triplex. A dsRNA-ssRNA junction structure can be pre-formed or induced by the binding of daPNAs (through the strand invasion of weakly formed dsRNA regions). We demonstrated the application of the new molecular recognition strategy in targeting a series of RNA structures. Dicer cleavage activity data suggest that daPNAs can be used to inhibit miRNA maturation in a substrate specific manner. Cell culture data show that daPNAs are effective in stimulating ribosomal frameshifting. Thus, the daPNAs platform would serve as programmable junction-specific molecular glues for the probing and targeting many biologically and medically essential RNA structures in transcriptomes.

## Methods

### Synthesis of PNA and RNA oligomers

Reverse-phase high-performance liquid chromatography (RP-HPLC) purified Cy3, Cy5 labeled and internal Cy3 labeled RNAs were purchased from Sigma-Aldrich and Biosyntech China, respectively. The PNA monomers thymine (T), cytosine (C), Guanine (G) and Adenine (A) were purchased from ASM Research Chemicals. PNA monomers L, Q, E, and S were synthesized following the reported methods(62-65). PNA oligomers were synthesized manually using Boc chemistry via a Solid-Phase Peptide Synthesis (SPPS) protocol. 4-Methylbenzhydrylamine hydrochloride (MBHA·HCl) polystyrene resins were used. The loading value used for synthesizing the oligomers was 0.3 mmol/g and acetic anhydride was used as the capping reagent. Benzotriazol-1-yl-oxytripyrrolidinophosphonium hexafluorophosphate (PyBOP) and *N,N*-diisopropylethylamine (DIPEA) were used as the coupling reagent. The oligomerization of PNA was monitored by the Kaiser test. Cleavage of the PNA oligomers was done using the trifluoroacetic acid (TFA) and trifluoromethanesulfonic acid (TFMSA) method, after which the oligomers were precipitated with diethyl ether, dissolved in deionized water and purified by reverse-phase high-performance liquid chromatography (RP-HPLC) using water-CH_3_CN-0.1% TFA as the mobile phase. LC-MS/MS were used to characterize the oligomers. The extinction coefficients of A, G, T, s^2^U, C, L, and Q were estimated to be 15.4, 11.7, 8.8, 10.2, 7.3, 7.3, and 7.3 mM^−1^cm^−1^, respectively, at 260 nm.

### Molecular modelling

We modelled the structure based on the major-groove PNA·RNA-RNA triplex structure from our previously studies modelled^2^. A three-base-pair PNA-RNA duplex segment was firstly extracted from a reported structure (PDB ID: 1PNN), then connected the PNA-RNA duplex segment to the remaining PNA·RNA-RNA triplex structure. Subsequently, the obtained structure was then solvated by TIP3P water molecules, followed by optimization, and NVT and NPT equilibrium. Then, a 100 ns product MD was conducted with the trajectory used for clustering. The centered structure of the most populated cluster was regarded as the representative structure. The structures were then all subject to minimization and NVT/NPT equilibrium, followed by a 30 ns simulated annealing (SA, 298-448-448-298 K). For the complexes with both PNA·RNA-RNA triplex and PNA-RNA duplex simultaneously formed, an extra 60 ns SA simulation was done with a higher temperature (298-498-498-298-498-498-298 K). During SA, all the base-base hydrogen bonds were restrained (0.0-0.3-0.4 nm, 1000 kJ/mol/nm^2^ for the distance restraint, and 180 degrees, 5.0 kJ/mol/rad^2^ for the angle restraints).

### Nondenaturing polyacrylamide gel electrophoresis

The nondenaturing polyacrylamide gel electrophoresis (PAGE) (20%, bis:arc = 29:1) experiments were performed using an incubation buffer containing 200mM NaCl, 0.5mM EDTA, 20mM HEPES, pH 7.5. For the labelled RNAs, the concentration was 5,10, 20 nM with varied PNA concentrations in 20 µL of buffer. Snap cooling was done for RNA hairpins by directly dipping the sample tubes from 95 °C into ice bath and annealing with PNAs by slow cooling from 65 °C to room temperature. The sample tubes were incubated for 2∼4 hours at 4 °C before the loading. To all the samples, 35% glycerol (20% of the total loaded volume) was added followed by vortexing for a few seconds before loading into the sample wells of the gels. Using a running buffer 1× TBE (Tris–Borate–EDTA), pH 8.3, the gels were run at 4 °C for 5 h with a voltage set at 250V for all the experiments. The gels were imaged using an Amersham Imager 680.

### Bio-layer interference assay

The 3′ biotin-RNA samples (Biosyntech) were diluted by binding buffer (containing 200mM NaCl, 0.5mM EDTA, 20mM HEPES, pH 7.5) into 200 nM final concentration. The concentrations of PNAs are varied for da5t (100, 50, 25, 12.5 and 6.25 nM), and for db198 and da198-AAA (4000, 2000, 1000, 500 and 250 nM). The association and dissociation time periods are both 300 seconds at 30 °C.

### *In vitro* dicing assay

The protocol for the *in vitro* dicing assay was described previously(66). Cy3 labelled pre-miR-198 and pre-miR-21 RNA oligomers synthesized by (Biosyntech) was used in the dicing assay. PNAs with varied concentrations were pre-incubated with RNA at 50°C for 10min, then slow cooling to room temperature (RT). After hDicer was added, the dicing system was incubated at 37 °C for 90min. The reaction was stopped with 2×RNA loading buffer containing 8 M Urea, 1×TBE, 0.05% Bromophenol blue, boiled for 10 min, and subsequently chilled on ice. RNA products were analyzed on 15% polyacrylamide, 8 M urea denaturing gel electrophoresis and visualized with a Typhoon Trio Imager (Amersham Biosciences).

### Plasmids, cloning, and mutagenesis

The programmed -1 ribosomal frameshifting (PRF) efficiency in a cell-free system was determined by p2luc plasmid (pDL for brevity)(49). As described previously, the model RNA (rHP1) served as an experiment construct and an in-frame control (IFC) plasmid were constructed by disrupting the slippery sequences, with a 100% PRF efficiency. To generate dual-fluorescence reporter (pDF) constructs, the sequences of rHP1-5m were inserted between the enhanced green fluorescent protein (eGFP) and monomer cherry protein (mCherry) coding sequences into the pcDNA5 vector BamHI and XhoI restriction enzyme site. The plasmids were extracted using Monarch Plasmid Miniprep kit (NEB) and precipitated by cold ethanol. The quality and concentration of plasmid DNA were measured by NanoDrop one spectrophotometer (Thermo Scientific). Plasmids with various mutations were generated by PCR methods using Q5^®^ Site-Directed Mutagenesis Kit (NEB) according to the instructions and verified by Sangon, China.

### DNA linearization and transcription

PCR was employed to prepare linearized DNA based on primers pDL-631 and pDL-3431 using Q5 Hot start PCR polymerase (NEB). All the components were mixed well and incubated in a thermal cycler (Biorad) under the following procedure: 95°C for 30 seconds; 35 cycles of 95°C for 30s, 57°C for 30s, 72°C 2 min; 72°C for 2min. The PCR products were precipitated by cold ethanol. The transcription process was driven by T7 polymerase using HiScribe T7 High Yield RNA Synthesis Kit (NEB). After 2h transcription, the DNA template was removed by DNase I (NEB) digestion and the RNA transcripts were purified by the RNA Transcript Purification Kit (NEB) and dissolved in RNase-free double-distilled H_2_O.

### *In vitro* translation

Translation protocols were described previously with minor modifications(26). In brief, 0.1 μM mRNA (final concentration) was mixed with various concentrations of PNAs and the appropriate volume of rabbit reticulocyte lysate (RRL) system (Promega). The translation mixtures were incubated at 30°C for 90 mins followed by the dual-luciferase reporter assay (Promega). The 2μl translation products were added into a white 384-well microplate (PerkinElmer), then 20 µl Dual-Glo reagent (Promega) and the relative luminescence units were measured using an Envision microplate reader (PerkinElmer) under the ultra-sensitive model. After the Fluc luminescence was measured, 20 μl Dual-Glo Stop reagent (Promega) was added into the above mixture to measure the *Rluc* luminescence. The PRF efficiency was calculated under the formula(49):

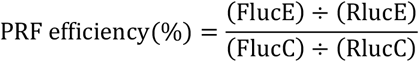

Here, FlucE and FlucC are the Fluc luminescence intensities of experimental and control samples, respectively. RlucE and RlucC are the Rluc luminescence intensities of experimental and control samples, respectively. Each experiment was performed at least thrice, and data were analyzed by GraphPad Prism 9.3 using the One-Way ANOVA method.

### Cell culture and PNA transfection

HEK-293T (CRL-3216) was a gift provided by Prof. Lei Yong. About 5×10^3^/well were seeded into 384-well cell culture plates, cultured in Dulbecco’s modified Eagle’s medium (Gibco) supplemented with 10% fetal bovine serum (BI), 100 units/ml of penicillin and 100 μg/ml streptomycin (Sangon) at 37°C in 5% CO_2_. The next day, 100 ng dual-fluorescent reporter plasmids with various concentrations of PNAs were co-transfected into cells using lipo3000 (Invitrogen) according to the instructions. After transfection for 24h, the eGFP and mCherry signals were imaged by Lionheart FXAutomated Microscope (Biotek) with a 4×objective under laser focus mode. The total intensities of eGFP and mCherry were transformed into grayscale and analyzed by CellProfiller Analyst with a threshold value of 0.15. The in-frame control plasmid with the slippery site disrupted was used as a control for calculating the PRF efficiency with the formula:

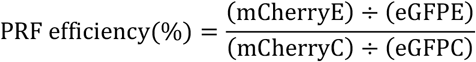

Here, mCherryE and mCherryC are the mCherry fluorescence intensities of experimental and control samples, respectively. eGFPE and eGFPC are the eGFP fluorescence intensities of experimental and control samples. Each experiment was performed at least three times, and data were analyzed by GraphPad Prism 9.3 using the One-Way ANOVA method.

## ASSOCIATED CONTENT

Supporting Information. MS data of List of DNA oligonucleotides used, PNA oligomers, nondenaturing PAGE assay data of PNA binding, cell-free dual luciferase reporter and cell culture dual fluorescent reporter assay data.

## AUTHOR INFORMATION

Corresponding Author

*Gang Chen

*Guobao Li

## Author Contributions

Gang Chen, Rongguang Lu, Guobao Li, Hongwei Wang and Zheng Liu conceived this project. Liping Deng, Ruiyu Lin, Yining Liu, Zhan Xuan, Alan Ann Lerk Ong, Manchugondanahalli S. Krishna, Kiran M. Patil, and Desiree-Faye Kaixin Toh synthesized the PNA oligomers. Liping Deng, Yun Lian, Yanyu Chen, and Alan Ann Lerk Ong performed the PAGE experiments. Rongguang Lu, Yun Lian, Yanyu Chen and Hanting Zhou performed the bio-layer interference assay. Rongguang Lu, Shijian Fan, Yanyu Chen, Hanting Zhou and Lixia Yang performed the cell-free dual-luciferase and cell culture dual-fluorescent assays. Xin Ke performed the Dicer enzyme cleavage activity assay. Kun Xi, Meng Zhenyu, Yunpeng Lu, Zheng Liu and Lizhe Zhu performed the molecular simulation study.

## Notes

The authors declare the following competing financial interest: A patent application based on some of the work reported here has been filed.

## ACKNOWLEDGMENTS

We thank Prof. Sam Butcher for sharing the experimental protocols, and for providing the p2luc plasmids, which was originally constructed by Prof. John Atkins and Raymond Gesteland. We thank Prof. Yongjuan Zhao and Dr. Yunjun Ge for their helpful suggestions on performing the dual-luciferase reporter assay. We thank Prof. Liang Yang for his help in using the Lionheart FX Automated Microscope (BioTek).

