## Supplementary material for "Recognition of RNA secondary structures with a programmable peptide nucleic acid-based platform": SI

### Table of Contents


| | | |
|---|---|---|
| 1 | Chemical structures of base triples: **Figure S1** | S3 |
| 2 | Mass spectrum and HPLC data of PNA oligomers: **Table S1 and Figures S2-S5** | S3-S5 |
| 3 | RNA and DNA oligomers used in this study: **Table S2** | S5 |
| 4 | Structural modelling data: **Figures S6 and S7** | S6 |
| 5 | Analysis of nondenaturing PAGE data for daPNA da5t and dbPNA P5 binding to rHP1-5t and other hairpins: **Figure S8** | S7 |
| 6 | Bio-layer interferometry assay data for daPNA da5t binding to rHP1 and its mutants: **Figure S9** | S7 |
| 7 | Plasmids used in this study and sequencing data: **Table S3 and Figure S10** | S8 |
| 8 | Normalized cell-free dual-luciferase, and cell culture dual-fluorescent protein assay data of da5t targeting rHP1-5m: **Figure S11** | S9 |
| 9 | Cell culture dual fluorescent protein reporter assay data for rHP1-5m and its mutants targeting with da5t: **Figure S12** | S10 |
| 10 | Nondenaturing PAGE data for da198-AAA binding to pre-miR-198: **Figure S13** | S10 |
| 11 | Bio-layer interferometry assay data for db198 binding to pre-miR-198: **Figure S14** | S11 |
| 12 | *In vitro* Dicer activity assay data of daPNA da198-AAA targeting pre-miR-21: **Figure S15** | S11 |
| 13 | Effect of da198-AAA on Dicer cleavage of pre-miR-21: **Figure S16** | S12 |
| 14 | Targeting tau pre-mRNA splice site hairpin structure by a daPNA: **Figure S17** | S12 |

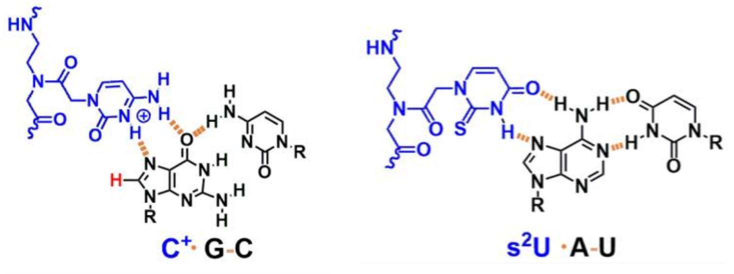


**Figure S1.** Chemical structures of base triples $C^{+}$·G-C and $s^{2}$U·A-U.

**Table S1.** Mass spectrum data for the synthesized PNAs.

| PNA | Sequence (N-terminus to C-terminus) | Molecular formula | Calculated MW | Observed MW |
|---|---|---|---|---|
| **da5t** | $NH_2$-Lys-AGAGTLTLTT-$CONH_2$ | $C_{114}H_{149}N_{57}O_{31}S_2$ | 2876.12 | 2876.12 |
| **da5t-NC** | $NH_2$-Lys-TGTATAQLL-$CONH_2$ | $C_{106}H_{144}N_{54}O_{27}S_2$ | 2669.10 | 2668.08 |
| **da198-AAA** | $NH_2$-Lys-LLTL2TLLAAA-$CONH_2$ | $C_{121}H_{159}N_{61}O_{28}S_6$ | 3106.12 | 3106.12 |
| **da-tau** | $NH_2$-Lys-TQTQTGCCG-$CONH_2$ | $C_{109}H_{157}N_{55}O_{31}$ | 2732.24 | 2728.22 |

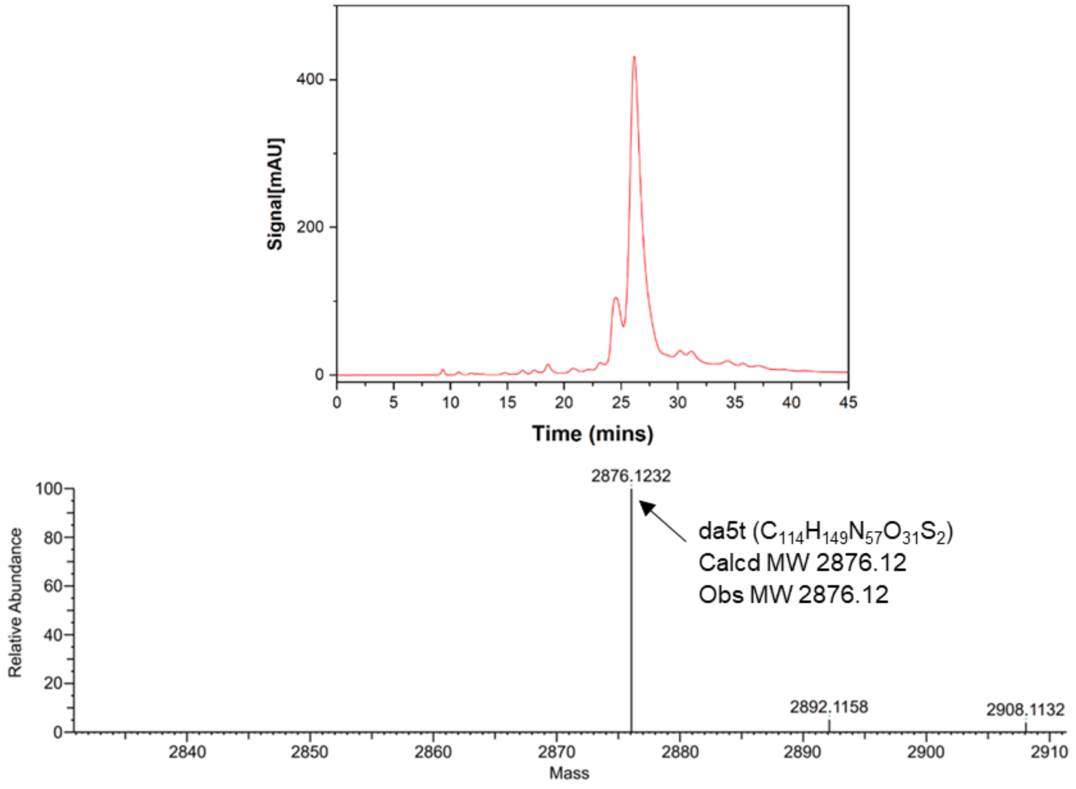


**Figure S2: HPLC (top) and LC/MS (bottom) data of da5t.**

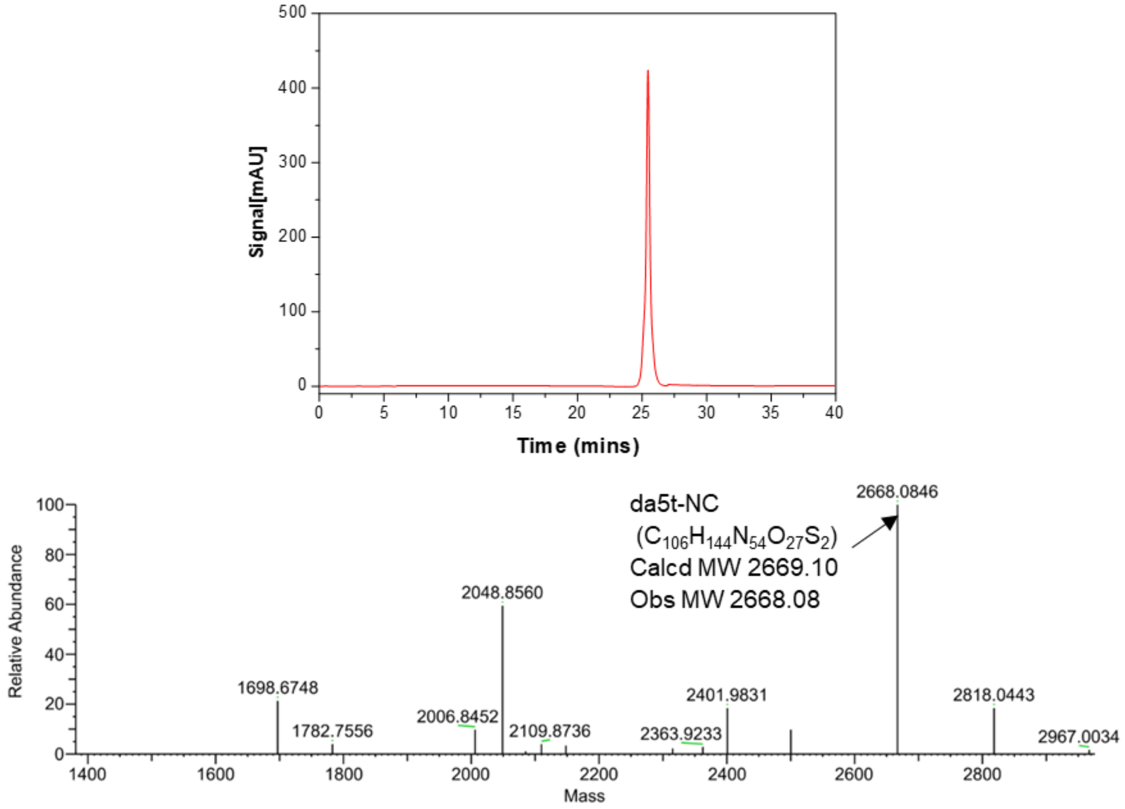


**Figure S3: HPLC (top) and LC/MS (bottom) data of da5t-NC.**

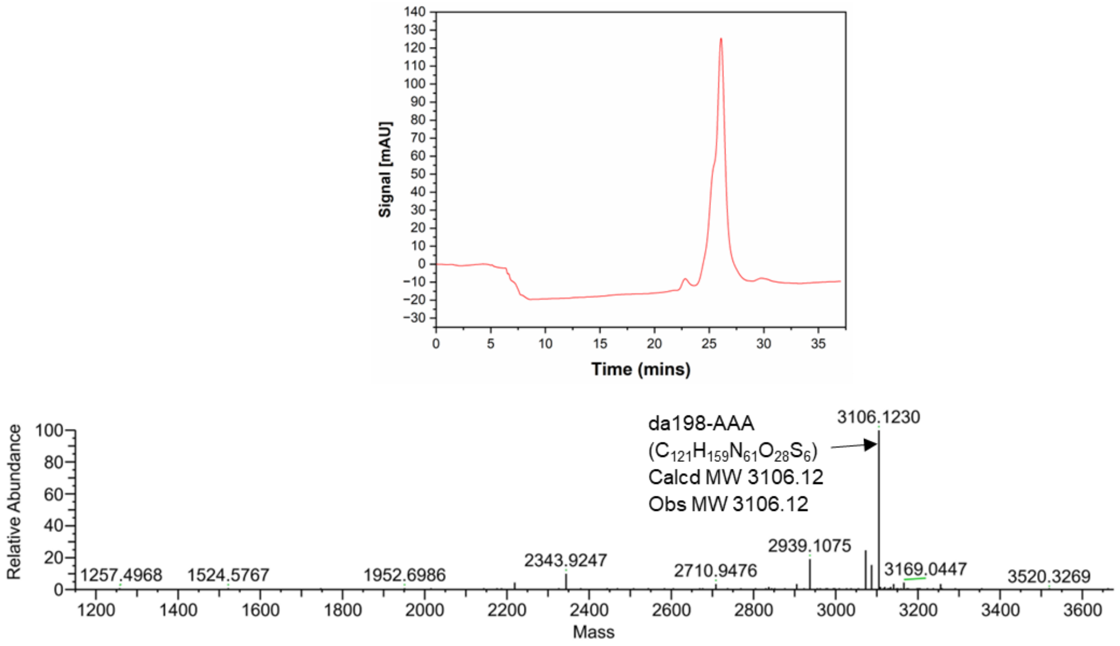


**Figure S4: HPLC (top) and LC/MS (bottom) data of da198-AAA.**

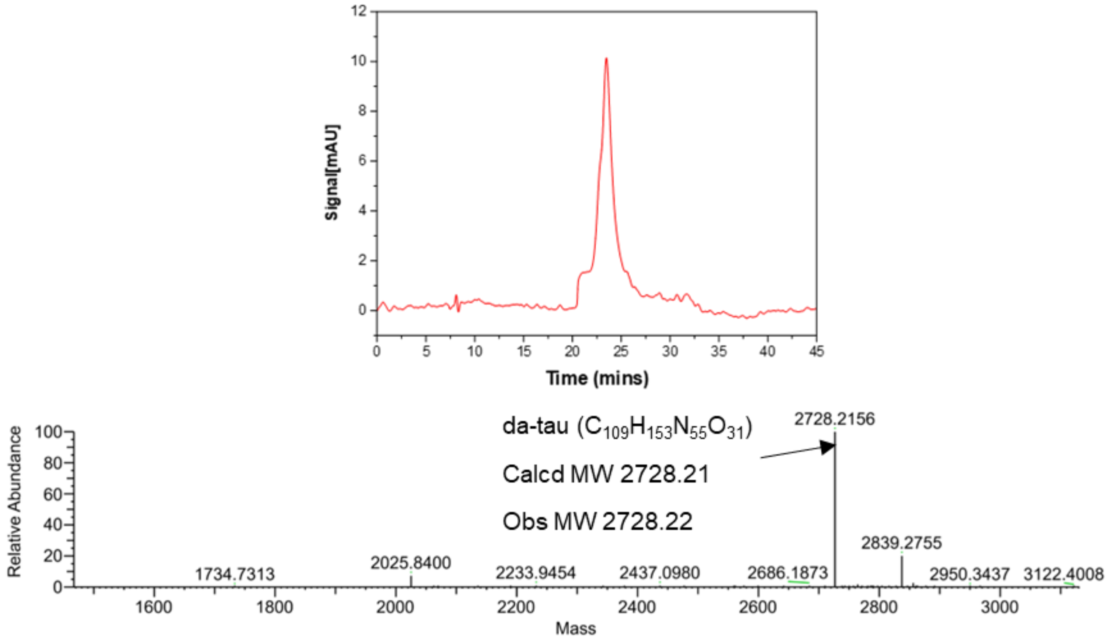


**Figure S5: HPLC (top) and LC/MS (bottom) data of da-tau.**

#### Table S2. RNA and DNA oligonucleotides used in this study.

| Oligomer | Label | Sequences (5'-3') | DNA/RNA |
|---|---|---|---|
| rHP1-iCy3 | Cy3 | UAGAGAGAGAAAGUU**[Cy3]**CGACUUUCUCUCUCUA | RNA |
| rHP1-5t-3′Cy5 | Cy5 | AGAGAAAGUUUCGACUUUCUCUCUCUA-**Cy5** | RNA |
| rHP1-5t-m1-3′Cy5 | Cy5 | AGAGAAAGUUUCGACUUUCUCUCcCUA-**Cy5** | RNA |
| rHP1-5t-m3-3′Cy5 | Cy5 | AcAGAAAGUUUCGACUUUCUgUCUCUA-**Cy5** | RNA |
| rHP1-5t-m4-3′Cy5 | Cy5 | cGAGAAAGUUUCGACUUUCUCgCUCUA-**Cy5** | RNA |
| rHP1-3t-3′Cy5 | Cy5 | AUCUCAGAGAAAGUUUCGACUUUCUCU-**Cy5** | RNA |
| rHP1-3′ biotin | biotin | UAGAGAGAGAAAGUUUCGACUUUCUCUCUCUA-**biotin** | RNA |
| rHP1-5t-3′ biotin | biotin | AGAGAAAGUUUCGACUUUCUCUCUCUA-**biotin** | RNA |
| rHP1-5t-m1-3′ biotin | biotin | AGAGAAAGUUUCGACUUUCUCUCcCUA-**biotin** | RNA |
| rHP1-5t-m3-3′ biotin | biotin | AcAGAAAGUUUCGACUUUCUgUCUCUA-**biotin** | RNA |
| rHP1-5t-m4-3′ biotin | biotin | cGAGAAAGUUUCGACUUUCUCgCUCUA-**biotin** | RNA |
| rHP1-3t-3′ biotin | biotin | AUCUCAGAGAAAGUUUCGACUUUCUCU-**biotin** | RNA |
| Pre-miR-21 | Cy3 | **Cy3**-UAGCUUAUCAGACUGAUGUUGACUGUUGAAUCUCAUGGC AACACCAGUCGAUGGGCUGU | RNA |
| Pre-miR-198 | Cy3 | **Cy3**-GGUCCAGAGGGGAGAUAGGUUCCUGUGAUUUUUCCUUCUUCUCU AUAGAAUAA | RNA |
| Pre-miR-198-m1 | Cy3 | **Cy3**-GGUCCAGAGGGGAGAUAGGUUCCUGUGAUaaccCCUUCUUCUCU AUAGAAUAA | RNA |
| Pre-miR-198-m2 | Cy3 | **Cy3**-GGUCCAGAGGGGAcAUAGGUUCCUGUGAUUUUUCCUUgUUCUCU AUAGAAUAA | RNA |
| Pre-miR-198-m3 | Cy3 | **Cy3**-GGUCCAGAGGGGAGAUAGcUUCCUGUGAUUUUUgCUUCUUCUCU AUAGAAUAA | RNA |
| Pre-miR-198 | biotin | GGUCCAGAGGGGAGAUAGGUUCCUGUGAUUUUUCCUUCUUCUCU AUAGAAUAA-**biotin** | RNA |
| pDL-631 | — | CACTCCCAGTTCAATTACAG | DNA |
| pDL-3431 | — | TCTTATCATGTCTGCTCGAA | DNA |

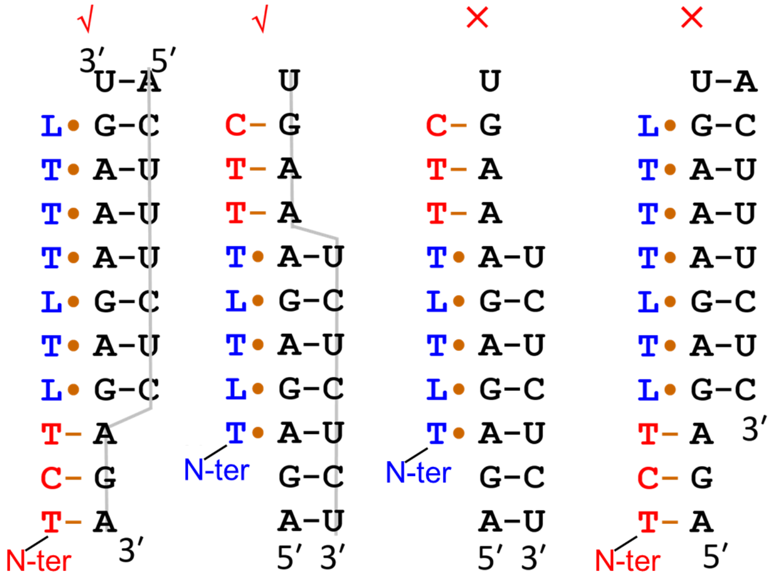

**Figure S6.** Modelling for daPNA targeting RNA structure. Only the first two complex structures form stably, with parallel dbPNA·dsRNA major-groove triplex and antiparallel asPNA-ssRNA duplex formed. The last two, however, show no PNA-RNA duplex structure formation, if the triplexes form.

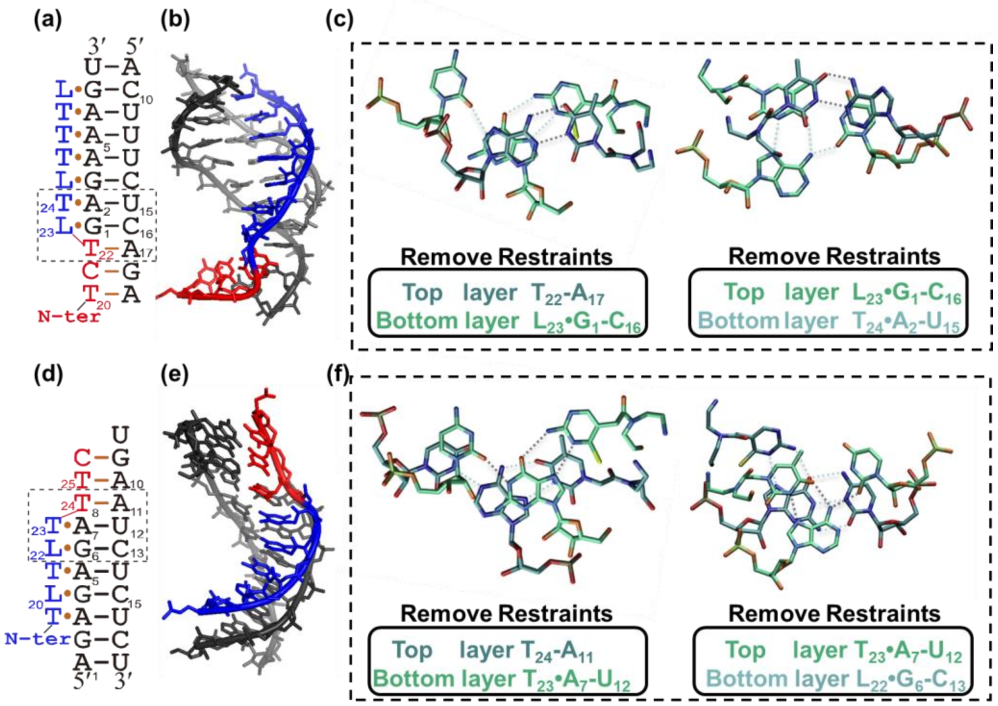


**Figure S7.** Testing the structural stability without base-base hydrogen bond restraints. (a, d) Schematic of the two binding modes for daPNA binding to RNA structure. (b, e) Molecular simulation models of daPNA binding to RNA structure without base-base hydrogen bond restraints. (c, f) Potential base pairs and base triples formed for the boxed regions in panels (a) and (d) in the absence of base-base hydrogen bond restraints.

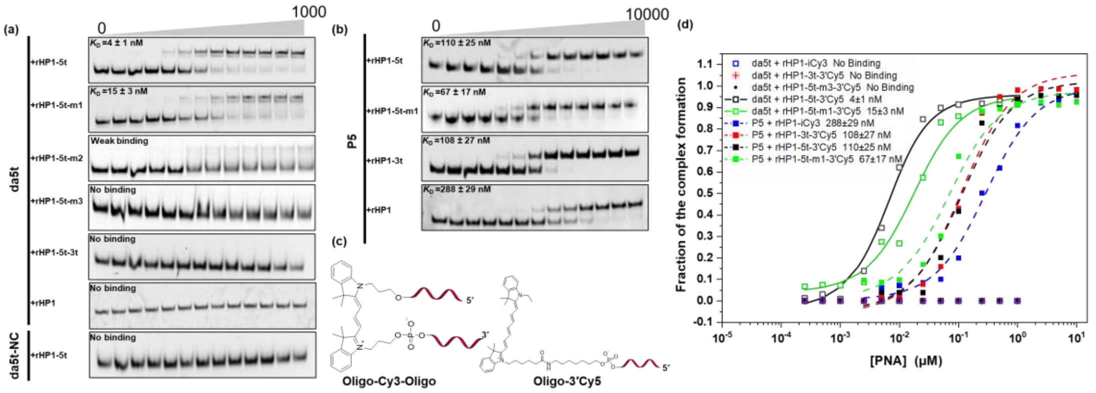

**Figure S8.** Nondenaturing PAGE data for da5t binding to rHP1-5t. (a) Binding data for daPNA da5t and da5t-NC. The RNA concentration loaded is 5 nM. The PNA concentrations in lanes from left to right are 0, 0.25, 0.5, 1, 2.5, 5, 10, 25, 50, 100, 250, 500, and 1000 nM, respectively. (b) Binding data for dbPNA P5. The RNA concentration loaded is 5 nM. The PNA concentrations in lanes from left to right are 0, 2.5, 5, 10, 25, 50, 100, 250, 500, 1000, 2500, 5000 and 10000 nM, respectively. (c) Chemical structures of Cy5 and Cy3 labelled RNAs. (d) The fitting curve of each PAGE results. The binding data for dbPNA P5 and daPNA da5t are shown with open symbol/dashed fitting line and solid symbol/solid fitting line, respectively.

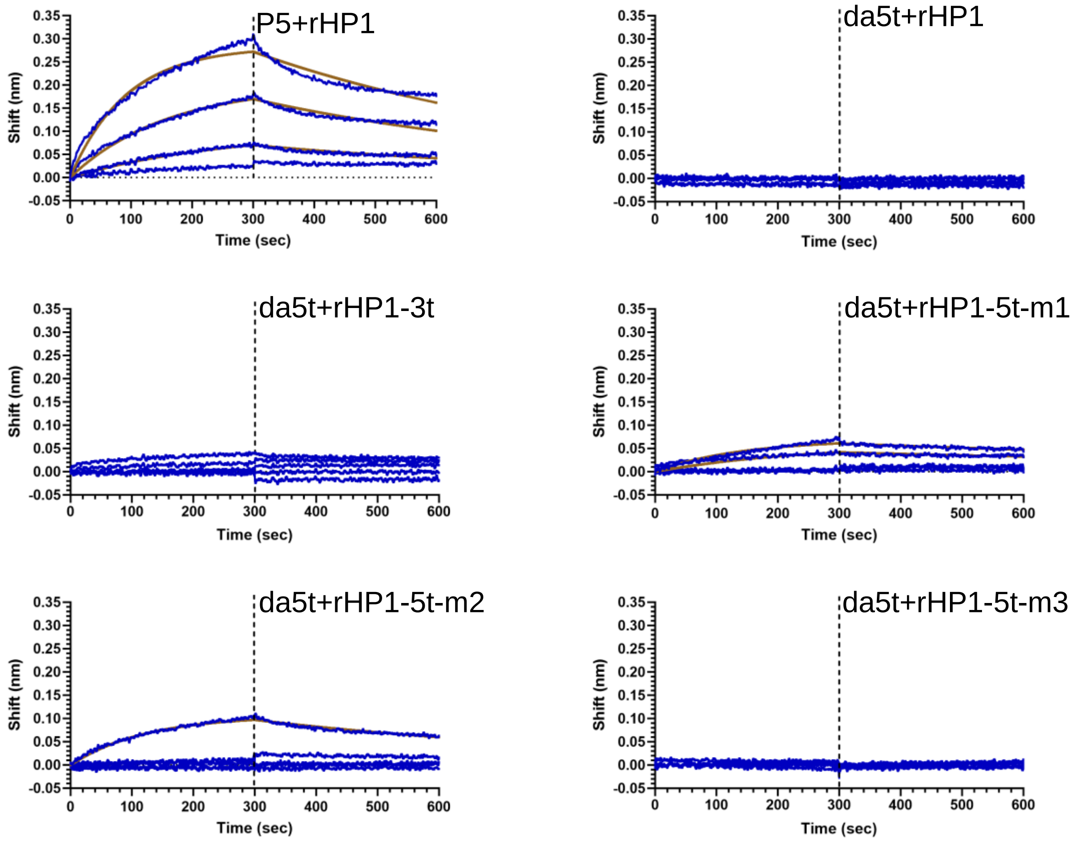


**Figure S9.** Bio-layer interferometry assay data for dbPNA P5 and daPNA da5t binding to rHP1 and its mutants. The experimental data and fitting curves are shown in blue and brown, respectively. The final concentrations of the dbPNA P5 are 20000, 10000, 5000 and 2500 nM from top to bottom. The final concentrations of the daPNA da5t are 100, 50, 25, 12.5, 6.25 nM from top to bottom.

#### Table S3. Plasmids used in this study.

| Plasmid | Vector | Inserted sequences (5'-3') |
|---|---|---|
| pDL-rHP1 | P2luc | TTTTTTAGGGTAGAGAGAGAAAGTTTCGACTTTCTCTCTCTA |
| pDL-rHP1-5m | P2luc | TTTTTTAGGGTgactAGAGAAAGTTTCGACTTTCTCTCTCTA |
| pDL-rHP1-5m-m1 | P2luc | TTTTTTAGGGTgactAGAGAAAGTTTCGACTTTCTCTCcCTA |
| pDL-rHP1-5m-m2 | P2luc | TTTTTTAGGGTgactcGAGAAAGTTTCGACTTTCTCgCTCTA |
| pDL-rHP1-5m-m3 | P2luc | TTTTTAGGGTgactAcAGAAAGTTTCGACTTTCTgTCTCTA |
| pDF-rHP1-5m | pcDNA5 | TTTTTTAGGGTgactAGAGAAAGTTTCGACTTTCTCTCTCTA |
| pDF-rHP1-5m-m1 | pcDNA5 | TTTTTTAGGGTgactAGAGAAAGTTTCGACTTTCTCTCcCTA |

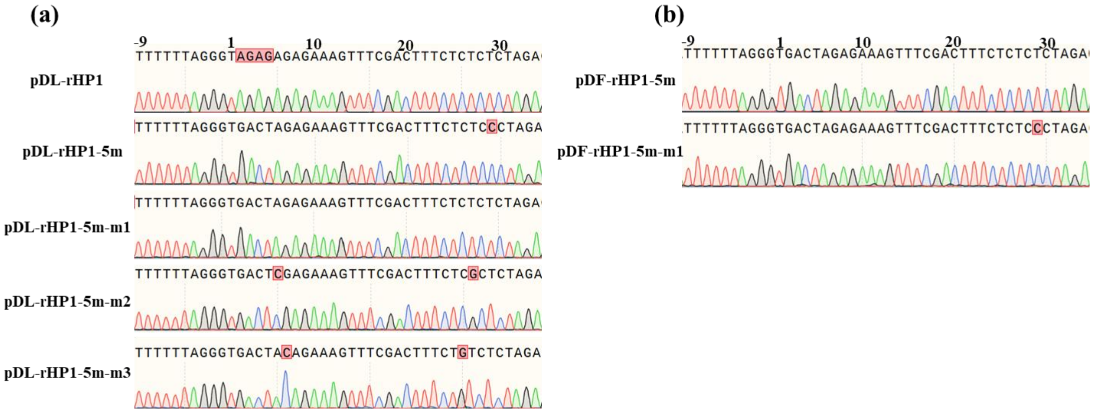


**Figure S10.** Sequencing data of plasmids. (a) Sequencing results of pDL plasmid rHP1 and its mutants involving pDL-rHP1, pDL-rHP1-5m, rHP1-5m-m1, rHP1-5m-m2 and rHP1-5m-m3. (b) Sequencing results of pDF plasmids involving pDF-rHP1-5m and pDF-rHP1-5m-m1. The plasmids were verified by Sanger method (Sangon).

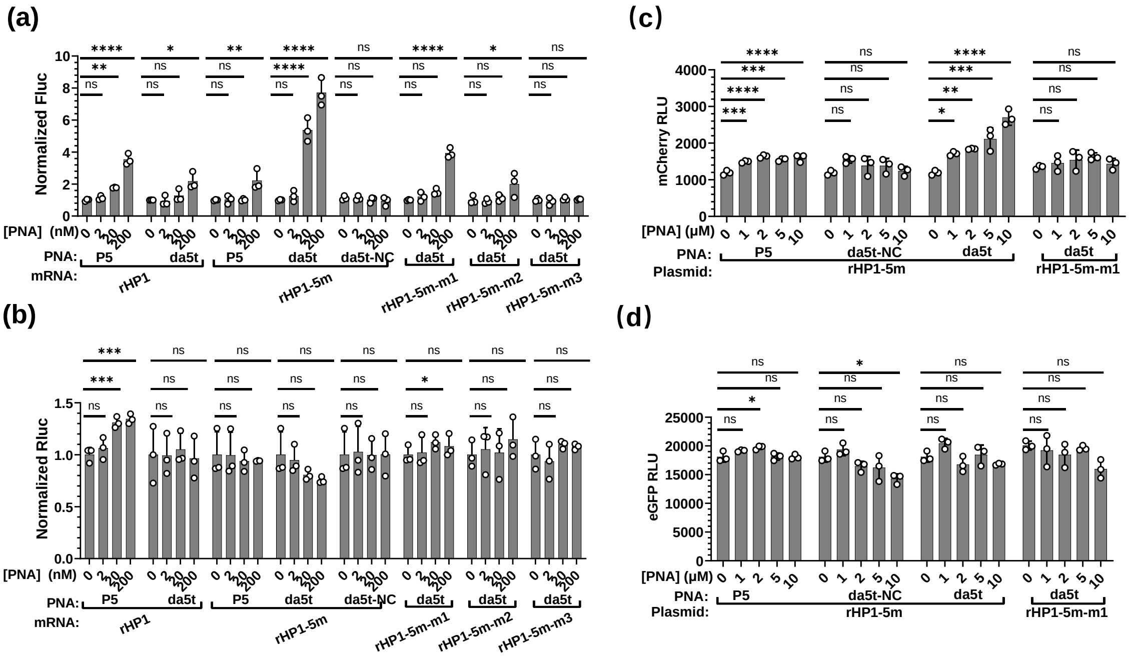


**Figure S11.** Normalized cell-free dual-luciferase, and cell culture dual-fluorescent protein assay data of da5t targeting rHP1-5m. (a, b) Normalized Fluc and Rluc activities in cell-free dual-luciferase reporter assay. (c, d) Normalized eGFP and mCherry levels in cell culture dual-fluorescent protein reporter assay. The data were analyzed by GraphPad Prism 9.3 and calculated by an ordinary one-way analysis of variance (ANOVA) using Dunnett's multiple comparisons test against the mean of mRNA or plasmid alone group. The error bars represent ± S.D. * $P < 0.05$, ** $P < 0.01$, *** $P < 0.001$, **** $P < 0.0001$, ns: not significant.

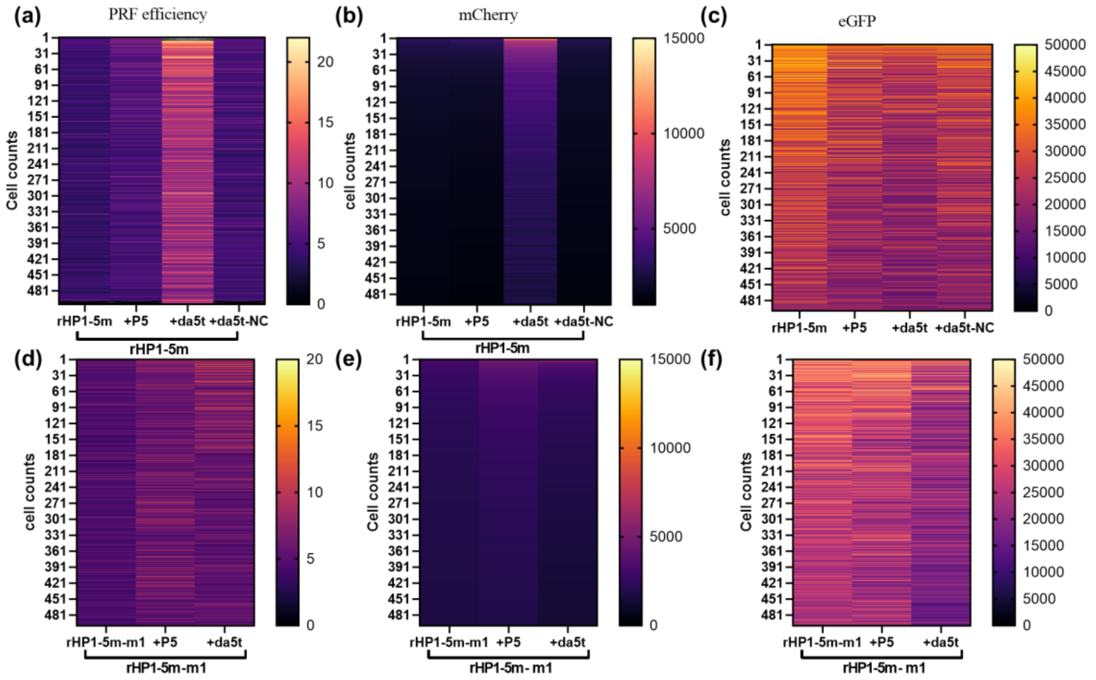


**Figure S12.** Cell culture dual fluorescent protein reporter assay data for rHP1-5m and its mutants targeting with da5t. Top 500 cells within each group were selected based on the mCherry RLU. (a-c) PRF efficiency value, mCherry and eGFP RLU of rHP1-5m targeting with dbPNA P5, daPNA da5t and control PNA da5t-NC. (d-f) PRF efficiency, mCherry and eGFP RLU of rHP1-5m-m1 targeting with dbPNA P5 and daPNA da5t. The final concentrations of PNA and plasmid are 10 μM and 4 ng/μl, respectively.

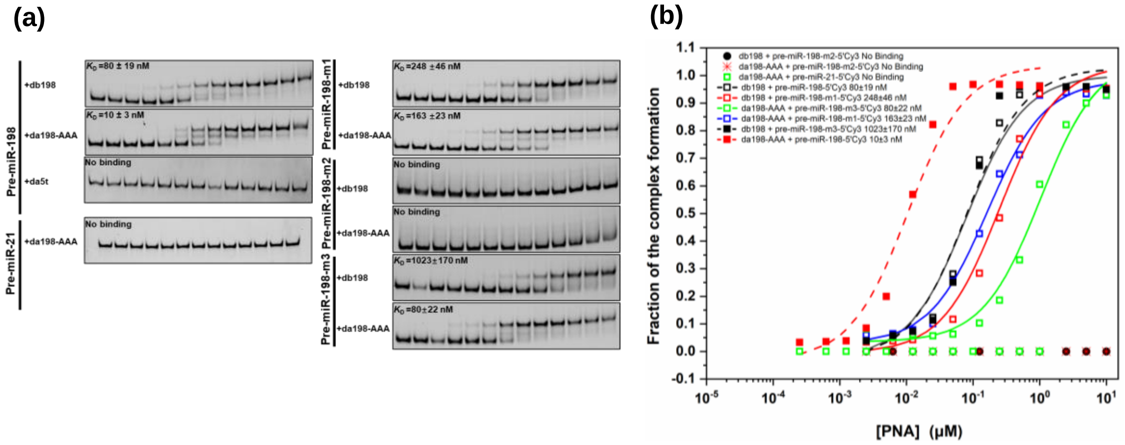


**Figure S13.** Nondenaturing PAGE data for db198 and da198-AAA binding to pre-miR-198. (a) Representative PAGE results. (b) Fitting curves of PAGE results. The RNA concentration loaded is 10 nM. The PNA concentrations in lanes from left to right are 0, 0.25, 0.5, 1, 2.5, 5, 10, 25, 50, 100, 250, 500, and 1000 nM, respectively.

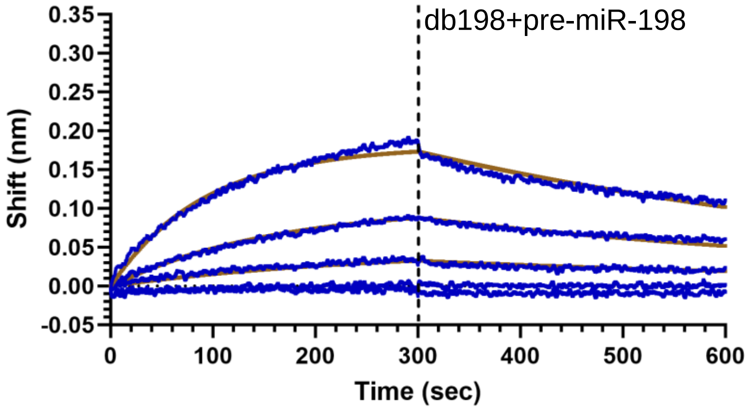


**Figure S14.** Bio-layer interferometry data for db198 and da198-AAA binding to pre-miR-198. The experimental data and fitting curves are shown in blue and brown, respectively. The final concentrations of the PNAs are 4000, 2000, 1000, 500, 250 nM from the top to bottom.

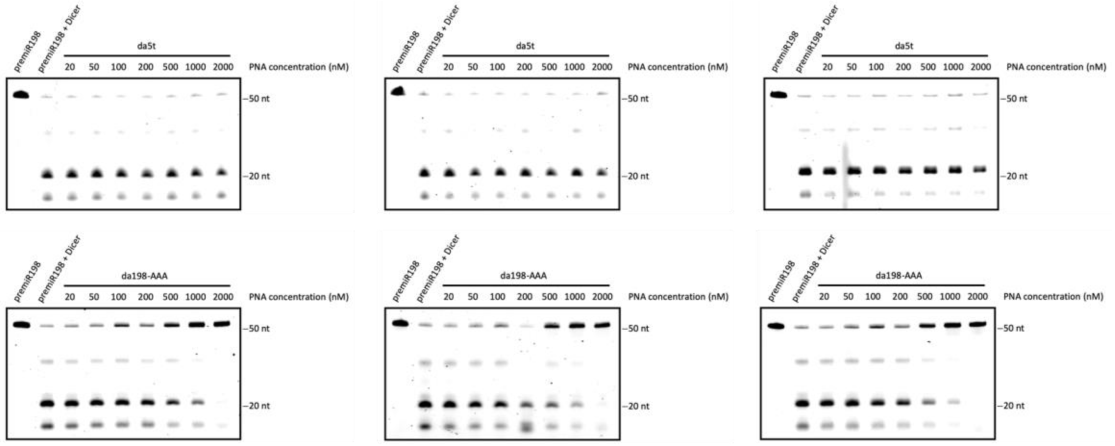


**Figure S15.** Effect of da198-AAA on Dicer cleavage of pre-miR-198. The PNA concentrations in lanes from left to right are 0, 20, 50, 100, 200, 500, 1000, 2000 nM, respectively.

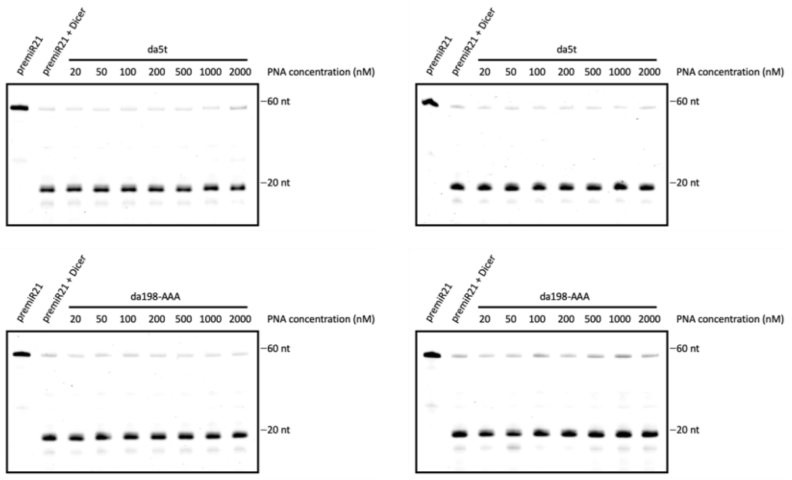


**Figure S16.** Test of the effect of da198-AAA on Decer cleavage of pre-miR-21. The PNA concentrations in lanes from left to right are 0, 20, 50, 100, 200, 500, 1000, 2000 nM, respectively.

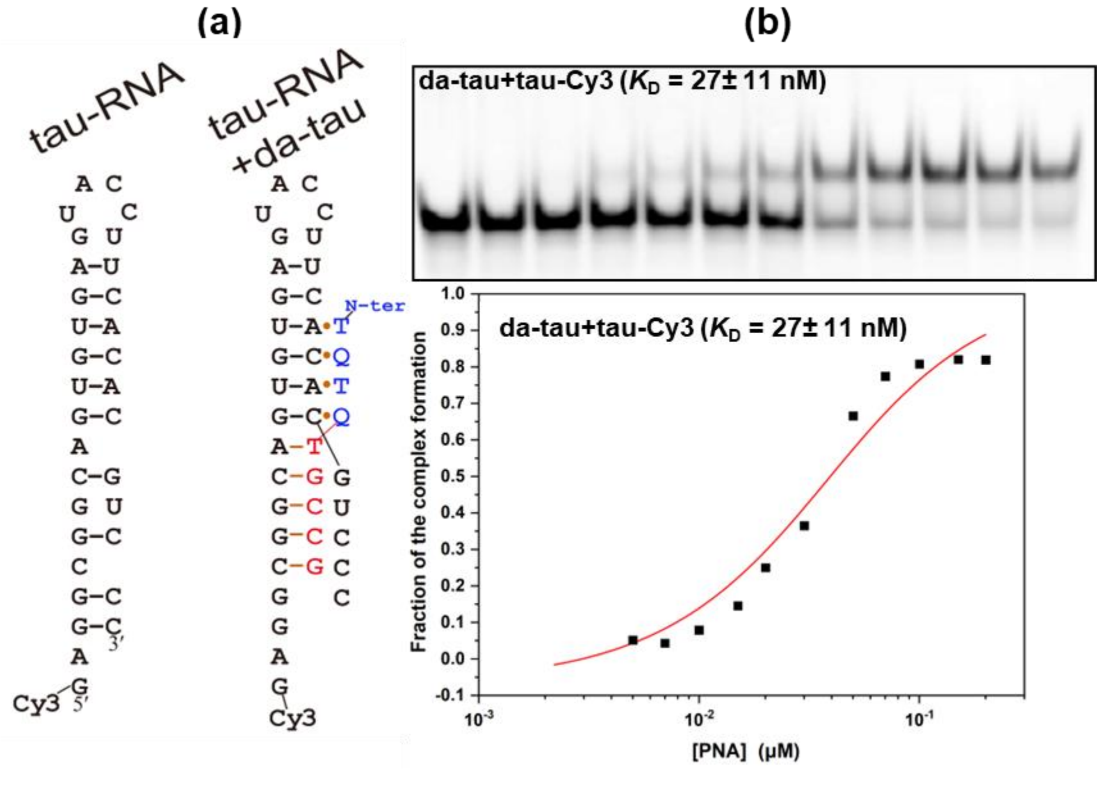


**Figure S17.** Targeting tau pre-mRNA splice site hairpin by daPNA. (**a**) Schematic of tau pre-mRNA hairpin construct and binding with PNAs. (**b**) Nondenaturing PAGE data and fitting curve of daPNA da-tau binding with tau pre-mRNA hairpin. The incubation buffer is 200 mM NaCl, 0.5 mM EDTA, 20 mM HEPES, pH 7.5. The running buffer is 1×TBE, pH 8.3. The loaded RNA hairpins are at 20 nM in 20 µL. The PNA concentrations in lanes from left to right are 0, 0.005, 0.007, 0.01, 0.015, 0.02, 0.03, 0.05, 0.07, 0.1, 0.15, and 0.2 µM, respectively. The gel shown is a representative gel. The error given is a standard error.
